# Evaluation of calcium-sensitive adenylyl cyclase AC1 and AC8 mRNA expression in the anterior cingulate cortex of mice with neuropathic pain

**DOI:** 10.1101/2021.09.29.462423

**Authors:** Stephanie Shiers, Hajira Elahi, Stephanie Hennen, Theodore J Price

## Abstract

The anterior cingulate cortex (ACC) is a critical region of the brain for the emotional and affective components of pain in rodents and humans. Hyperactivity in this region has been observed in neuropathic pain states in both patients and animal models and ablation of this region from cingulotomy, or inhibition with genetics or pharmacology can diminish pain and anxiety. Two adenylyl cyclases (AC), AC1 and AC8 play an important role in regulating nociception and anxiety-like behaviors through an action in the ACC, as genetic and pharmacological targeting of these enzymes reduces mechanical hypersensitivity and anxiety-like behavior, respectively. However, the distribution of these ACs in the ACC has not been studied in the context of neuropathic pain. To address this gap in knowledge, we conducted RNAscope *in situ* hybridization to assess AC1 and AC8 mRNA distribution in mice with spared nerve injury (SNI). Given the key role of AC1 in nociception in neuropathic, inflammatory and visceral pain animal models, we hypothesized that AC1 would be upregulated in the ACC of mice following nerve injury. This hypothesis was also founded on data showing increased AC1 expression in the ACC of mice with zymosan-induced visceral inflammation. We found that AC1 and AC8 are widely expressed in many regions of the mouse brain including the hippocampus, ACC, medial prefrontal cortex and midbrain regions, but AC1 is more highly expressed. Contrary to our hypothesis, SNI causes an increase in AC8 mRNA expression in NMDAR-2B (Nr2b) positive neurons in the contralateral ACC but does not affect AC1 mRNA expression. Our findings show that changes in *Adcy1* mRNA expression in the ACC are insufficient to explain the important role of this AC in mechanical hypersensitivity in mice following nerve injury and suggest a potential unappreciated role of AC8 in regulation of ACC synaptic changes after nerve injury.

## Introduction

Adenylyl cyclases (ACs) catalyze the formation of cyclic AMP (cAMP) which is a key signaling molecule regulating the activity of protein kinase A (PKA) (Kandel, 2012). In addition to other actions such as phosphorylation of transcription factors, PKA plays an important role in regulating neuronal excitability by phosphorylating voltage gated ion channels, and it also regulates synaptic plasticity by coordinating trafficking and channel kinetics of ionotropic glutamate receptors including AMPA and NMDA receptors (Kandel, 2012; Wang and Zhang, 2012). There are 10 ACs (*Adcy* gene family) in most mammalian genomes (Kandel, 2012). Nine of these are membrane-bound ACs with 12 transmembrane segments ACs (1-9) and one is a soluble AC (sAC), each of these enzymes exhibits different regulatory properties and expression patterns (Sadana and Dessauer, 2009). Canonical activation of these ACs involves stimulation of a G protein-coupled receptor that signals via a Gα_s_ subunit to bind and activate an AC enzyme. AC1 and AC8, which are predominantly expressed in the CNS, are known to be stimulated by Ca^2+^, in a calmodulin-dependent manner, and are commonly referred to as Ca^2+^-sensitive ACs (Xia and Storm, 1997; Wang and Zhang, 2012). In line with this neuron-specific expression, these ACs play specialized roles in synaptic plasticity in the spinal cord and brain of rodents (Wang et al., 2011; Griggs et al., 2019) where they augment NMDAR currents upstream of PKA phosphorylation of these channels (Liauw et al., 2005; Wang and Zhang, 2012). Genetic knockout of AC1 in mice demonstrates an important role for the *Adcy1* gene in chronic hypernociception that occurs after nerve injury or other injuries (Vadakkan et al., 2006; Xu et al., 2008; Wang et al., 2011; Corder et al., 2013; Griggs et al., 2019; Zhou et al., 2021). Genetic knockout of *Adcy8* suggests a specialized role for this enzyme in anxiety-like behavior in mice (Bernabucci and Zhuo, 2016) but does not support a role in mechanical hypersensitivity after inflammation, although formalin nocifensive behavioral responses are attenuated in mice lacking the *Adcy8* gene (Wei et al., 2002).

AC1 promotes mechanical hypersensitivity via an action in the spinal cord and in the anterior cingulate cortex (ACC) of mice (Liu et al., 2020) and plays a critical role in inflammatory, muscle, visceral and neuropathic injury models, and in promotion of latent nociceptive sensitization (Vadakkan et al., 2006; Xu et al., 2008; Wang et al., 2011; Wang and Zhang, 2012; Corder et al., 2013; Qiu et al., 2014; Brust et al., 2017; Griggs et al., 2019; Liu et al., 2020; Zhou et al., 2021). Small molecule inhibitors of AC1 have been developed, such as NB-001 and ST034307, and these molecules reverse mechanical hypersensitivity behaviors in mice and rats, consistent with the notion that AC1 promotes pain through synaptic plasticity mechanisms in the CNS (Wang et al., 2011; Brust et al., 2017; Cheng et al., 2019; Griggs et al., 2019; Zhou et al., 2021). Direct injection of NB-001 into the ACC inhibits mechanical hypersensitivity in multiple mouse pain models in both sexes (Wang et al., 2011; Liu et al., 2020; Zhou et al., 2021), suggesting that AC1 activity in the ACC is a key site for promotion of persistent pain. The ACC has long been recognized as an important brain region for the affective component of pain (Talbot et al., 1991; Rainville et al., 1997; Hofbauer et al., 2001). This is further supported by the fact that anterior cingulotomy has been recognized as a surgical option for the management of intractable cancer and non-cancer pain in humans (Hassenbusch et al., 1990; Wilkinson et al., 1999). Collectively, these lines of evidence makes AC1 an attractive potential target for pain therapeutics.

While Ca^2+^-sensitive ACs, encoded by *Adcy1 and Adcy8,* are recognized as important mediators of chronic changes in nociception in animal models (Wang et al., 2011; Wang and Zhang, 2012), their mRNA expression distribution in the frontal cortex has not been examined in the context of a neuropathic injury model in male and female mice. In these studies, we used RNAscope *in situ* hybridization to define changes in *Adcy1* and *Adcy8* expression in cellular populations in frontal cortical areas in the spared nerve injury (SNI) neuropathic pain model in mice. We find that *Adcy1* is widely expressed in important brain regions for pain, where it colocalizes with neurons that express NMDA receptors, but only *Adcy8* shows increased expression in the cingulate cortex in the SNI model. Our work raises the testable hypothesis that AC8 may contribute to anxiety in neuropathic pain.

## Methods

### Animals

All animal procedures were approved by the University of Texas at Dallas Institutional Animal Care and Use Committee. Female and male C57BL/6 mice were used for these experiments. Mice were 4 weeks of age at the start of experiments. For assessment of expression of markers across the mouse brain (experiment 1), a single male mouse was used. For the sham versus SNI mouse expression comparison (experiment 2), 12 mice were used (6 sham and 6 SNI) with 3 males and 3 females per group.

### Surgery

Spared nerve injury was performed by the ligation and cutting of the tibial and peroneal branches of the left sciatic nerve trifurcation, leaving the sural branch intact. Sham surgeries were performed the same way but without ligating/cutting the nerve. A single injection of the antibiotic gentamicin (3mg/kg s.q.) was given immediately following surgery. Mice were allowed to recover for 2 weeks following surgery before being assessed for neuropathy.

### Mechanical withdrawal threshold

Tactile sensitivity was measured by probing the left outer surface of the left hind paw with a series of calibrated von Frey filaments. Withdrawal thresholds were calculated using the up-down method (Chaplan et al., 1994). Mechanical withdrawal thresholds were assessed in all animals before and after (2 weeks) SNI to confirm neuropathic pain.

### Tissue preparation

Mice were decapitated under isoflurane and the brain was removed and frozen in powdered dry ice. The frozen brain was slowly embedded in OCT by adding thin layers of OCT around the sample in order to avoid tissue thawing. The OCT blocks were then stored in a −80°C freezer. On the day of the experiment, the tissues were removed from the −80°C, placed on dry ice, and transferred to the −20°C cryostat chamber for ~30 minutes for temperature acclimation. The brain was then sectioned coronally at 20 μm onto charged slides. For assessment of markers across the mouse brain, single sections of main regions throughout the brain were kept including the medial prefrontal cortex-prelimbic cortex (mPFC - prelimbic), hippocampus (CA1, CA2, CA3), dentate gyrus (DG), lateral amygdala (LA), basolateral amygdala (BLA), mediodorsal thalamus, medial habenula, septal nuclei, caudate putamen, lateral septum, piriform cortex, periaqueductal gray (PAG), retrosplenial cortex, and anterior cingulate cortex (Cg1, Cg2). These regions were located by following a mouse brain atlas (Paxinos and Franklin, 2004) and looking for anatomical landmarks on the specimen.

For the sham versus SNI mouse experiments, at 3 weeks post-SNI, the animals were euthanized and the brains were prepared as described above. 20 μm sections targeting Cg1, Cg2 anterior cingulate cortex (Bregma +1.1 mm to −0.22 mm) were kept. While sectioning, a hole was made using a needle through the lateral portion of the hemisphere contralateral to injury. The slides were stored in a −80°C freezer.

### RNAscope and Nissl Staining

On the day of RNAscope experiments, the slide box was removed from the −80°C freezer, placed on dry ice, and transferred to the lab. The slides were immediately immersed in cold 10% formalin (pH 7.4) and processed for RNAscope using the fresh frozen protocol with a 1–2-minute Protease IV digestion as described by Advanced Cell Diagnostics (ACD; acdbio.com). The probes used are shown in **Table 1**. The combinations used are shown in individual images. The DAPI step was not performed on all sections. Instead, following completion of RNAscope, the slides were submerged in a coplin jar containing cold blocking solution (10% Normal Goat Serum, 0.3% Triton-X 100 in 0.1M Phosphate Buffer (PB)) for 1 hour at room temperature. The coplin jar was covered in tin foil to protect the slides from light-exposure. Following blocking, the slides were rinsed with 0.1M PB. Each slide was then placed in light-tight humidity control tray and Blue Fluorescent Nissl stain (Molecular Probes Neurotrace 435/455; Invitrogen Cat N21479) diluted in blocking solution (1:250) was pipetted onto the slide within the hydrophobic boundaries. This was performed one slide at a time to prevent tissue drying. The slides were incubated for 30 minutes, and then washed in 0.1M PB. The slides were then air dried, cover slipped with Prolong Gold and allowed to cure over-night.

**Table 1.** RNAscope probes.

| mRNA | ACD Probe Cat No. |
| --- | --- |
| *Adcy1* | 451241-C2 |
| *Adcy8* | 462041 |
| *Nr2a (Grin2a)* | 481831 |
| *Nr2b (Grin2b)* | 417391-C3 |

### Imaging and Analysis

Images were acquired on an Olympus FV3000 confocal microscope at 20X magnification. For experiment 1, a single image was acquired of all major brain regions listed above. For experiment 2, one 20X image was acquired of the contralateral Cg1 and Cg2 from each section, and 3-5 sections were imaged per animal (n=6 per group; 3 males and 3 females). The acquisition parameters were set based on guidelines for the FV3000 provided by Olympus (gain = 1, HV ≤ 600, offset = 4) and only laser power was adjusted. Given that probe intensity varies depending on the number of bound amplifiers (thereby, fluorescence intensity is not representative of abundance in this technology), each image was acquired using optimal settings to best visualize the mRNA puncta as instructed by ACD. Images were brightened/contrasted and analyzed in Olympus CellSens (v1.18).

To estimate mRNA abundance in experiment 2, ~25 *Nr2b, Adcy1,* and *Adcy8* co-positive neurons were randomly selected across all laminae in each image and their somas (Nissl signal) were traced using the polygon ROI tool. The area of *Adcy1* and *Adcy8* signal within the ROI was measured using the Count and Measure tool which highlights the mRNA signal using a thresholded detection. A manual threshold was applied to each image so that all mRNA signal was highlighted within the ROI. Any signal detected that was smaller than 1 μm was automatically not highlighted by the program. Since mRNA puncta is estimated to be ~1.5μm^2^ in size, we divided the area of detected *Adyc1* and *Adcy8* signal by 1.5 to estimate the number of mRNA puncta. The number of puncta was then divided by the area of the ROI (neuronal soma) to calculate *Adcy1* puncta/μm^2^ or *Adcy8* puncta/μm^2^.

### Statistics

Statistical tests are described in figure legends. An α value of p < 0.05 was considered significant.

## Results

We first assessed expression of *Adcy1, Adcy8, Nr2a* (also known as N-Methyl D-Aspartate Receptor Subtype 2A; GluN2A; Grin2a)*, Nr2b* (also known as N-Methyl D-Aspartate Receptor Subtype 2B; GluN2B; *Grin2b*) in a single male mouse. Our probe combinations in this set of experiments were *Adcy1/Nr2a/Nr2b* and *Adcy1/Adcy8.* We chose *Nr2a* and *Nr2b* as these NMDAR subunits predominate in the forebrain and are important downstream targets of AC activity (Liauw et al., 2005). We compared marker expression to the *in situ* hybridization database on the Allen Brain Atlas (ABA) and saw similar probe-specificity in all assessed regions. For instance, in the hippocampus, *Adcy1* showed low expression in the CA1 and CA3 subregions but was more pronounced in CA2 and the DG (**Fig 1A-B**) while *Adcy8* had very low expression in CA1 (**Fig 1B**). This pattern closely resembled the ABA expression map (**Fig 2C**) supporting that our probes were target-specific. We saw widespread expression of *Adcy1* throughout the brain and limited *Adcy8* expression (**Fig 1-2**; **S Fig 1-4**). *Adcy1* was highly expressed in the cortical areas of the brain including the dorsal (Cg1) and ventral (Cg2) areas of the mouse ACC, the prelimbic cortex, retrosplenial cortex c and amygdala, and appeared to coexpress *Nr2a* and *Nr2b* (**Fig 2; S Fig 1**). *Adcy1* showed little expression in the caudate putamen, medial habenula, medio-dorsal thalamus, and PAG (**S Fig 1-2**). *Adcy1* showed higher expression in all brain regions examined than *Adcy8* (**S Fig 3-4**). These results largely parallel Allen Brain Atlas (mouse.brain-map.org) and mousebrain.org datasets (Zeisel et al., 2018). Additionally, data from the proteinatlas.org shows that AC1 protein expression is higher in the human brain than AC8 but both are enriched in brain versus other tissues (Thul et al., 2017).

**Figure 1.**
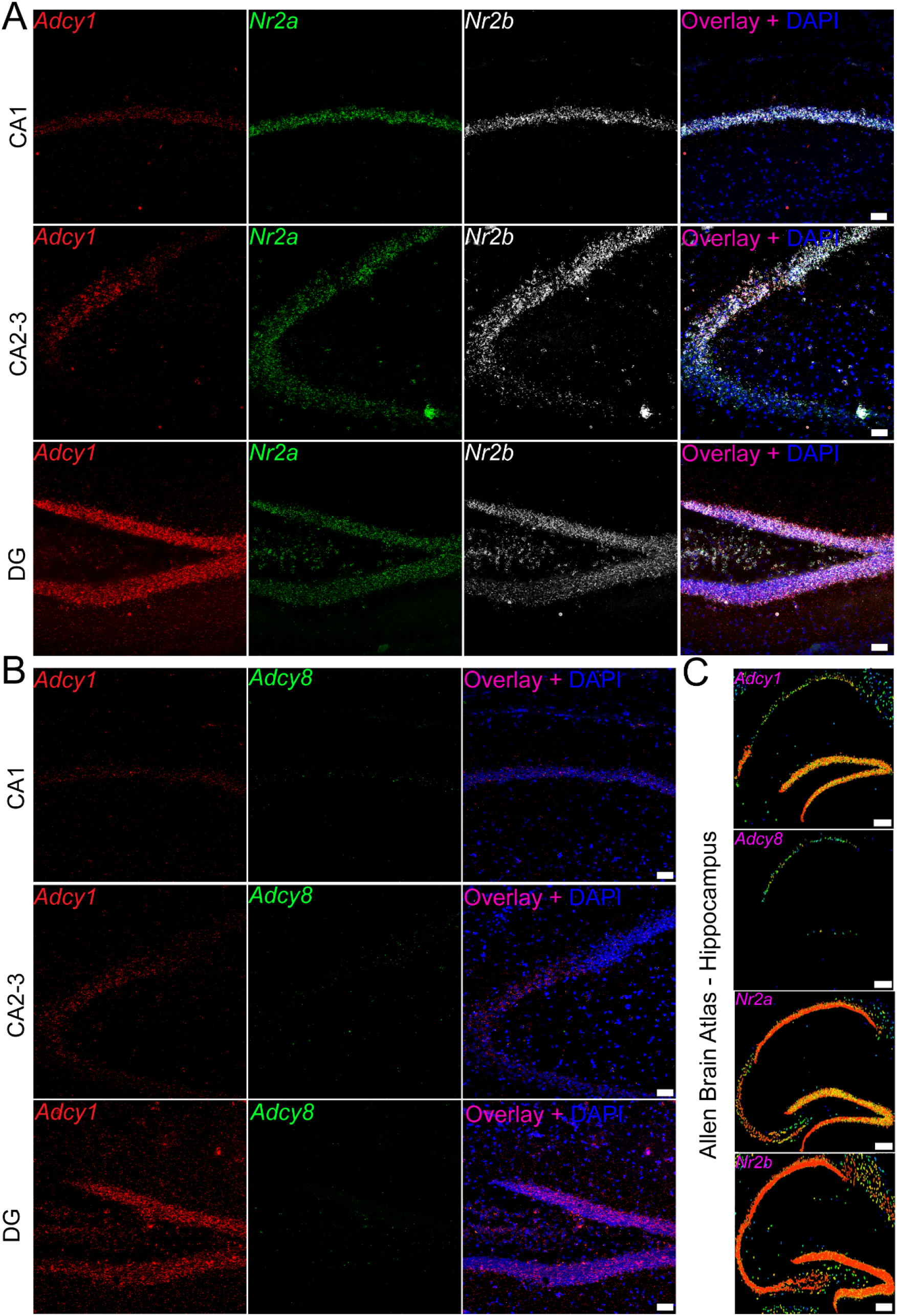
Example of RNAscope *Adcy1, Adcy8, Nr2a,* and *Nr2b* probe specificity in the brain. We conduced RNAscope *in situ* hybridization for *Adcy1/Nr2a/Nr2b* and *Adcy1/Adcy8* on a single mouse brain. To confirm probe specificity, we compared expression of these markers to the *in situ* hybridization database on the Allen Brain Atlas. **A)** For example, in the CA1, CA2 and dentate gyrus (DG) subregions of the hippocampus, *Adcy1* (red), *Nr2a* (green), *Nr2b* (white) and **B)** *Adcy1* (red), and *Adcy8* (green) show differential expression in abundance and regionspecificity **C)** which closely resembles the expression map found on the Allen Brain Atlas. We confirmed there was similar expression in all other assessed regions as well. All other images can be found in supplementary figures 1-4. Panel A-B: Images are 20X; scale bar = 50 μm. Panel C: scale bar = 210 μm.

**Figure 2.**
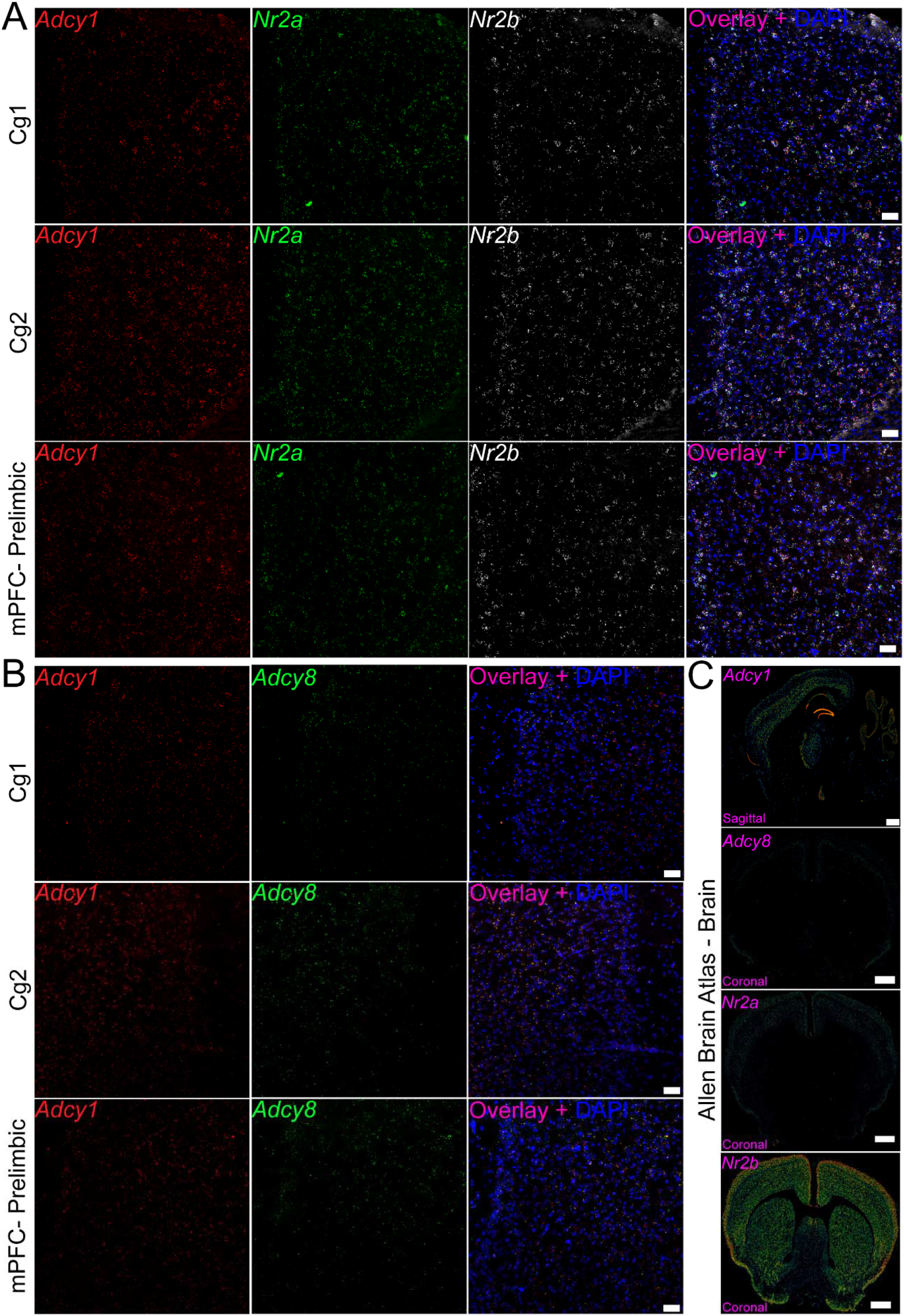
*Adcy1, Adcy8, Nr2a,* and *Nr2b* probe expression in the frontal cortex. We conduced RNAscope *in situ* hybridization for *Adcy1/Nr2a/Nr2b* and *Adcy1/Adcy8* on a single mouse brain. **A)** *Adcy1* (red), *Nr2a* (green), *Nr2b* (white) and **B)** *Adcy1* (red), and *Adcy8* (green) in the dorsal anterior cingulate cortex (Cg1), ventral anterior cingulate cortex (Cg2) and medial prefrontal cortex (mPFC)-prelimbic area. *Adcy1* had higher expression in the cortex than *Adcy8* **C)** which closely resembles the expression map found on the Allen Brain Atlas (coronal sections include Cg1/Cg2). We assessed coronal images of the ABA expression map when available. We confirmed there was similar expression in all other assessed regions as well. All other images can be found in supplementary figures 1-4. Panel A-B: Images are 20X; scale bar = 50 μm. Panel C: scale bar = 932 μm.

After establishing the specificity of these probes, and the overlap of *Adcy1* with *Adcy8, Nr2a* and *Nr2b* expression in many brain regions, we sought to determine if *Adcy1* or *Adcy8* mRNA expression was altered in the SNI model of neuropathic pain. SNI surgery was conducted, and the presence of mechanical hypersensitivity was established by von Frey testing 3 weeks after surgery. SNI mice showed mechanical hypersensitivity compared to sham controls (**S Fig 5**). Brains were taken from mice and then processed for mRNA assessment. We examined the ACC because AC1 has been associated with neuropathic pain-like behaviors through a mechanism that involves the ACC in mice (Xu et al., 2008; Wang et al., 2011; Brust et al., 2017; Zhou et al., 2021). To determine changes in mRNA expression we estimated signal abundance in neurons as shown in **Fig 3**. We used a Nissl staining technique to label whole neurons, and then calculated the area of the neuron covered with mRNA signal for each probe to estimate abundance (**Fig 3A-C**). We then used this method to assess changes in *Adcy1* and *Adcy8* mRNA abundance in neurons of the dorsal (Cg1) and ventral (Cg2) areas of the mouse ACC on the side contralateral from SNI or sham surgery. We found an increase in *Adcy8* abundance in neurons of the Cg1 area, but no change in *Adcy1* mRNA (**Fig 4**). In the Cg2 region neither *Adcy1* nor *Adcy8* were significantly increased by SNI (**Fig 5**). We used an equal split of male and female animals in these experiments. No sex differences were noted, but the sample size was not large enough to directly assess sex differences.

**Figure 3.**
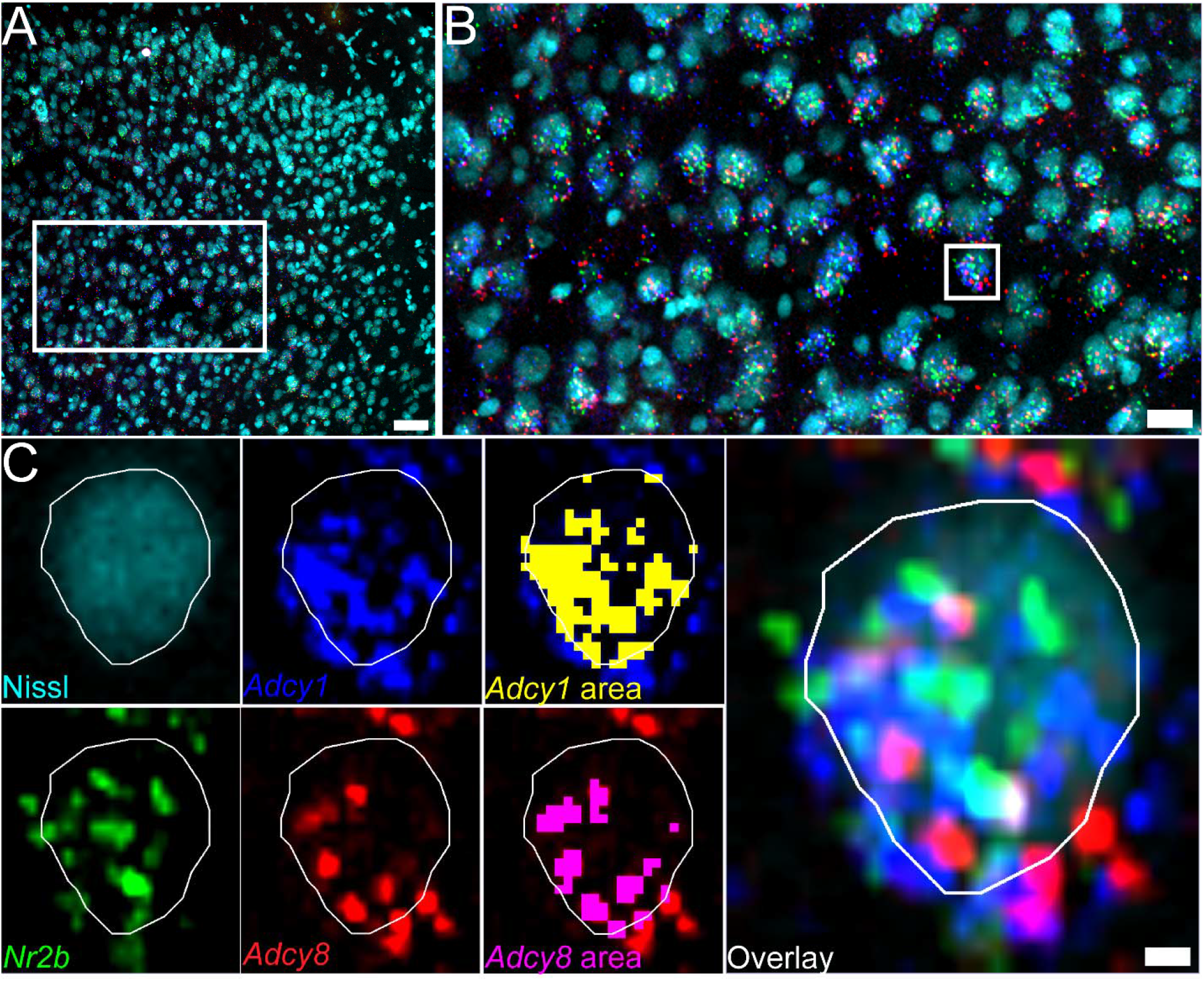
mRNA abundance analysis method. **A)** Example image of Nissl (cyan), *Nr2b* (green), *Adcy1* (blue), and *Adcy8* (red) signal in the Cg1 region of the anterior cingulate cortex. 20x image; Scale bar = 50 μm. **B)** Zoom-in of area highlighted (white box) in panel A. Scale bar = 20 μm. **C)** Example of how the abundance analysis is conducted on a single neuron (white box shown in Panel B). The soma of a randomly selected *Nr2b, Adcy1, Adcy8* co-positive neuron is traced using the Nissl stain as boundaries. The area of *Adcy1* and *Adcy8* signal is detected using the Count and Measure tool in Olympus CellSens. The area is then divided by 1.5 μm^2^, the estimated average size of an mRNA puncta, to approximate mRNA puncta / μm^2^. Scale bar = 2 μm.

**Figure 4.**
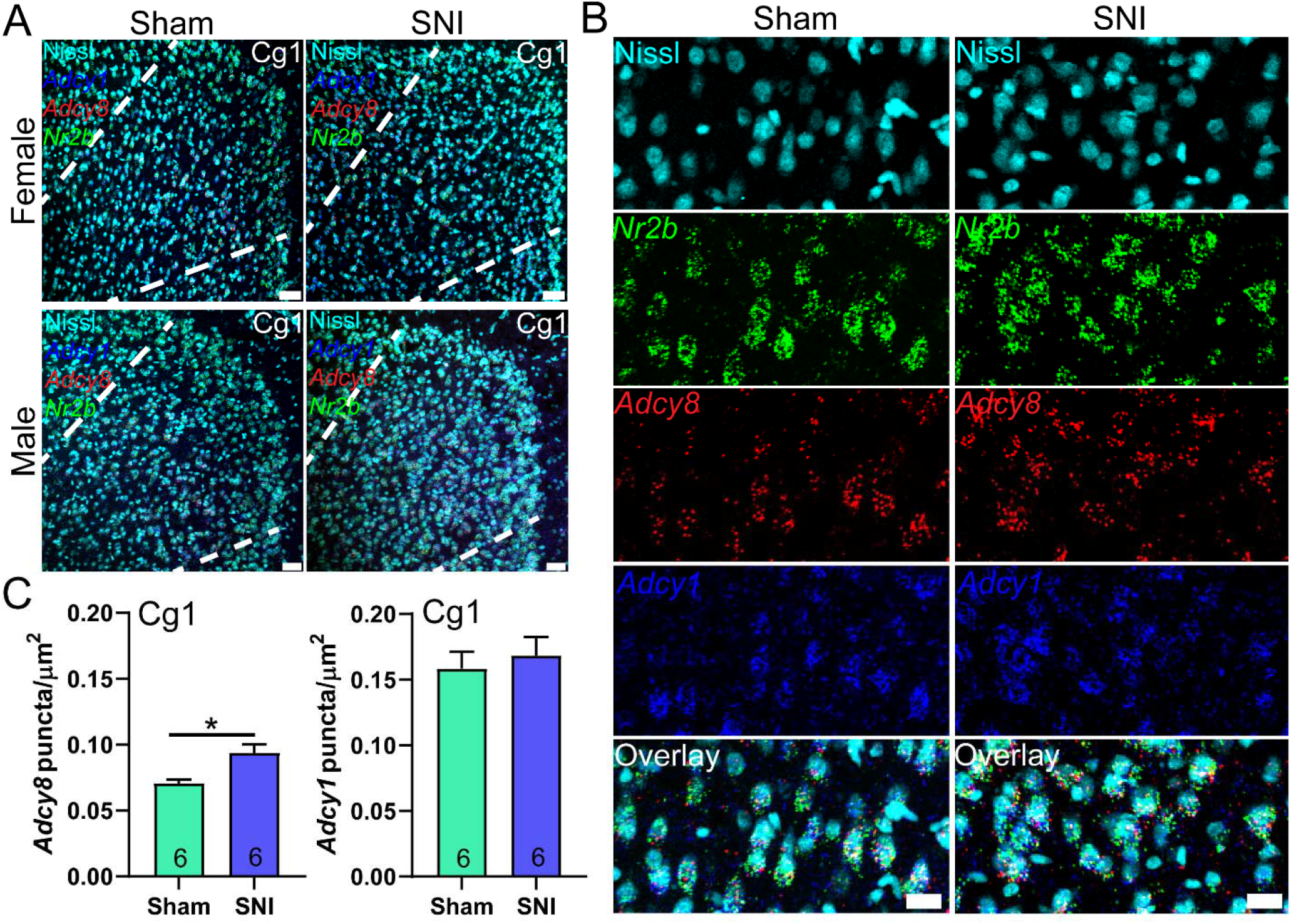
*Adcy1* and *Adcy8* mRNA abundance in the dorsal anterior cingulate cortex (Cg1). **A)** 20X images of Cg1 (area between white dotted lines) stained for *Adcy1* (blue), *Adcy8* (red), *Nr2b* (green) and Nissl (cyan) from male and female mice with SNI or sham surgeries. Scale bar = 50 μm. **B)** A cropped zoomed-in image shows mRNA signal in Cg1 neurons. Scale bar = 20 μm **C)** Significantly more *Adcy8* mRNA puncta was detected in *Nr2b+* neurons in Cg1 of SNI mice. Unpaired t-test *p<0.05.

**Figure 5.**
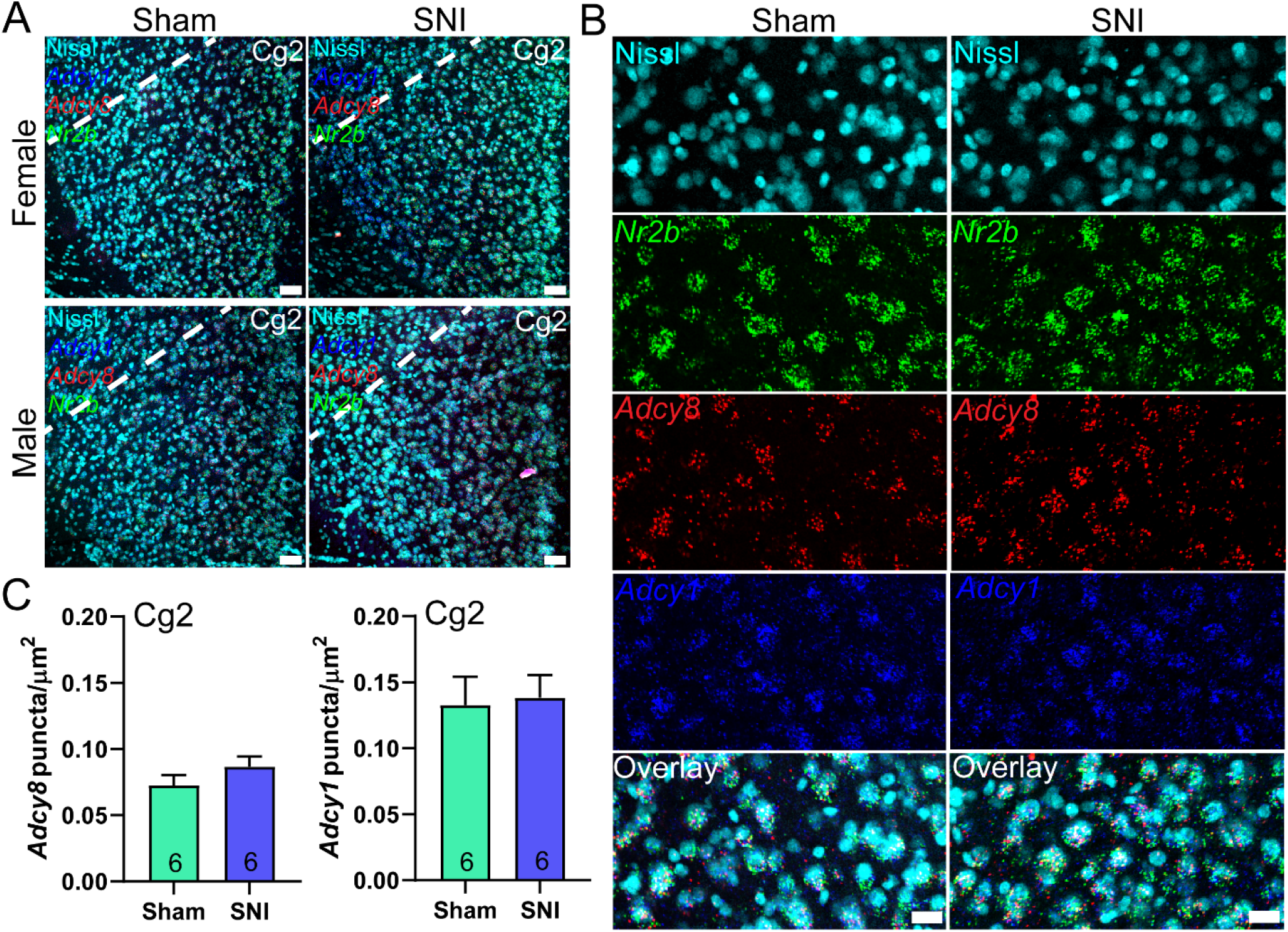
*Adcy1* and *Adcy8* mRNA abundance in the ventral anterior cingulate cortex (Cg2). **A)** 20X images of Cg2 (area below white dotted line) stained for *Adcy1* (blue), *Adcy8* (red), *Nr2b* (green) and Nissl (cyan) from male and female mice with SNI or sham surgeries. Scale bar = 50 μm. **B)** A cropped zoomed-in image shows mRNA signal in Cg2 neurons. Scale bar = 20 μm. **C)** There was no significant difference in amount of *Adcy1* nor *Adcy8* mRNA puncta / μm^2^ in the Cg2 subregion between SNI and sham mice. Unpaired t-test p>0.05.

## Discussion

Our findings confirm that AC1 and AC8 are expressed throughout the mouse brain including many cortical regions. AC1 is more widely expressed, corroborating previous brain-wide expression mapping studies done by the Allen Brain Institute (mouse.brain-map.org) and the Human Protein Atlas (proteinatlas.org). Given the extensive literature on the ACC’s involvement in the affective aspects of pain in people (Hassenbusch et al., 1990; Talbot et al., 1991; Rainville et al., 1997; Wilkinson et al., 1999; Hofbauer et al., 2001) and its involvement in chronic pain models in rodents as well as the role of AC1/AC8 in modulating injury-associated and anxiety-like behaviors in this region (Xu et al., 2008; Blom et al., 2014; Pereira et al., 2014; Gu et al., 2015; Kang et al., 2015; Sharim and Pouratian, 2016; Sellmeijer et al., 2018; Deng et al., 2019), we investigated whether AC1 and AC8 mRNA expression in this region changed after nerve injury. In the SNI mouse model of neuropathic pain, 3 weeks after injury, we found that *Adcy8* mRNA expression was increased in the dorsal, but not ventral ACC, likely reflecting an increase in *Adcy8* expression in neurons that either did not express *Adcy8* previously, or only expressed it on a low level. Based on previous experiments demonstrating increased AC1 expression in the zymosan visceral inflammation model in mice (Liu et al., 2020), we expected to observe increased *Adcy1* expression in the dorsal ACC, but our findings did not support our original hypothesis. Our findings do not discount that AC1 plays an important role in neuropathic pain-like behaviors in mice as both knockout of *Adcy1* gene, and inhibition of AC1 with pharmacological blockers, reduces neuropathic mechanical hypersensitivity in mice (Wei et al., 2002; Wang et al., 2011).

A hypothesis emerging from our findings that could be tested in future experiments is that increased expression of Adcy8 in the ACC may be linked to anxiety produced by neuropathic pain. The presence of neuropathic pain in humans often causes comorbid anxiety (Nicholson and Verma, 2004). While anxiety caused by peripheral nerve injury has not been consistently observed in mouse models (LaGraize et al., 2004; Urban et al., 2011; Sheahan et al., 2017), some studies suggest that mice and rats can develop anxiety after nerve injury (Seminowicz et al., 2009; Sellmeijer et al., 2018; Li et al., 2021), or develop changes in gene expression consistent with anxiety phenotypes (Descalzi et al., 2017). Previous studies using *Adcy8* knockout mice suggest that AC8 may be important for persistent anxiety caused by environmental cues (Bernabucci and Zhuo, 2016). A possible interaction of the *Adcy8* gene with SNI-induced anxiety could be tested in future studies.

There are several limitations to our study. We were not powered to look at sex differences, but we did use mice of both sexes. Given the small experimental variability, we think our findings of increased *Adcy8* expression in the Cg1 region likely occurs in both sexes. We only examined a single time point after nerve injury, but we chose a time point that is consistent with most published studies in the field. Anxiety may develop at later time points after injury (Seminowicz et al., 2009), so we may expect greater changes in *Adcy8* as time goes on. We did not examine potential changes in other brain regions because the existing literature has focused mostly on Ca^2+^-sensitive ACs in the ACC in pain models. Finally, we did not clarify the cell types in the ACC that show changes in *Adcy8* expression, but these neurons clearly express the NMDAR subunit, NR2B.

We conclude that the extensive literature on the role for AC1 in enhanced nociception after injury in mice cannot be explained by increased expression of Adcy1 mRNA in neurons in the ACC. This does not discount the possibility that protein levels may increase or that enzyme activity might increase due to enhanced Ca^2+^ signaling in these neurons after peripheral nerve injury. Nevertheless, our work highlights a potentially unappreciated role for AC8 that can be further explored using transgenic mice or through pharmacological manipulation. The latter may require further development of tool compounds as most pharmacological development on Ca^2+^-sensitive ACs has focused on AC1.

## Notes

**Conflict of Interests:** SS, HE and TJP declare no conflicts of interest. SH is an employee of Grünenthal GmbH.

### Competing Interest Statement

SS, HE and TJP declare no conflicts of interest. SH is an employee of Grunenthal GmbH.

