## Supplementary Figures for "Evaluation of calcium-sensitive adenylyl cyclase AC1 and AC8 mRNA expression in the anterior cingulate cortex of mice with neuropathic pain"

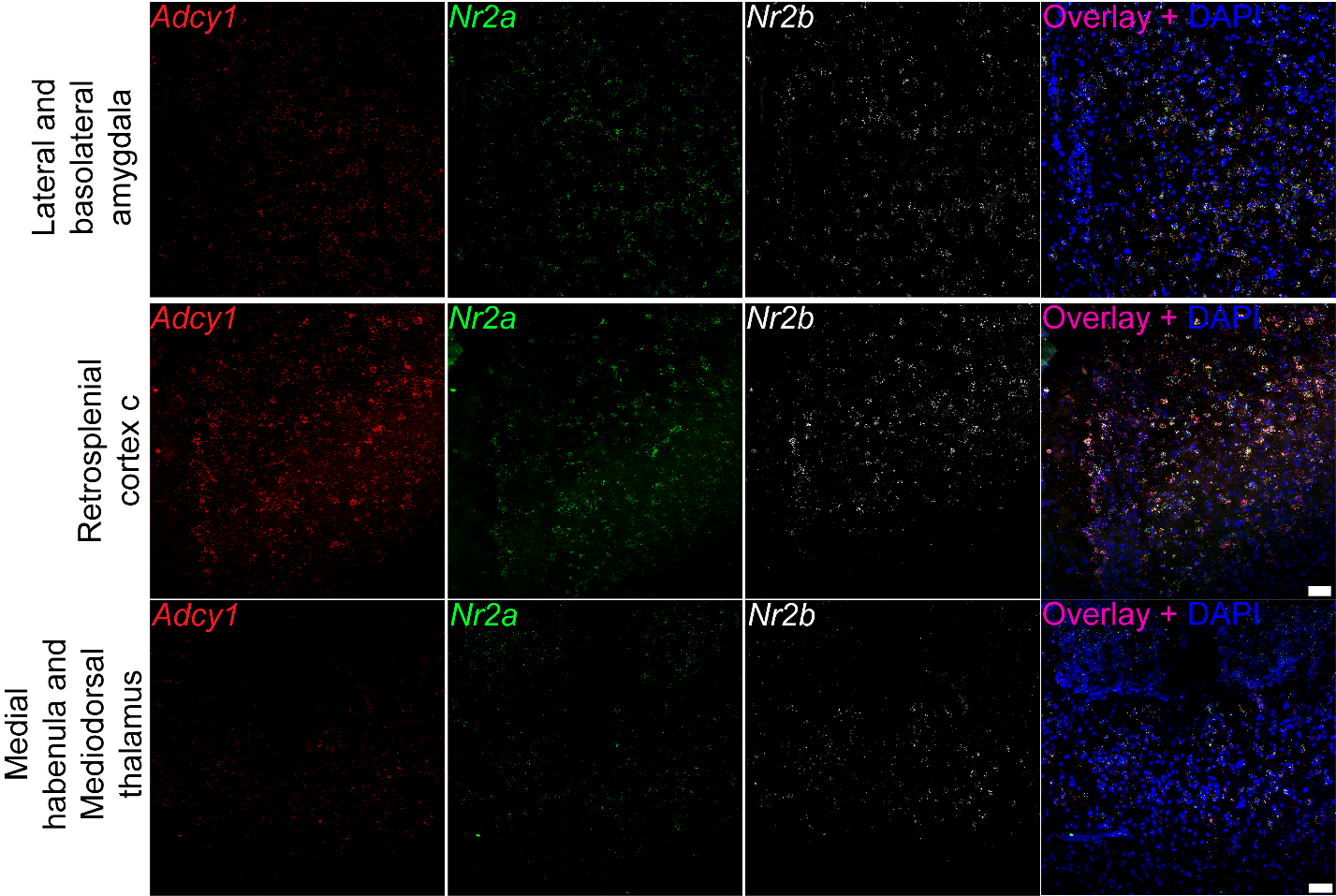

**Supplementary Figure 1. *Adcy1*, *Nr2a*, and *Nr2b* in the lateral and basolateral amygdala, retrosplenial cortex c, medial habenula, mediodorsal thalamus.** 20x; Scale bar = 50 µm.

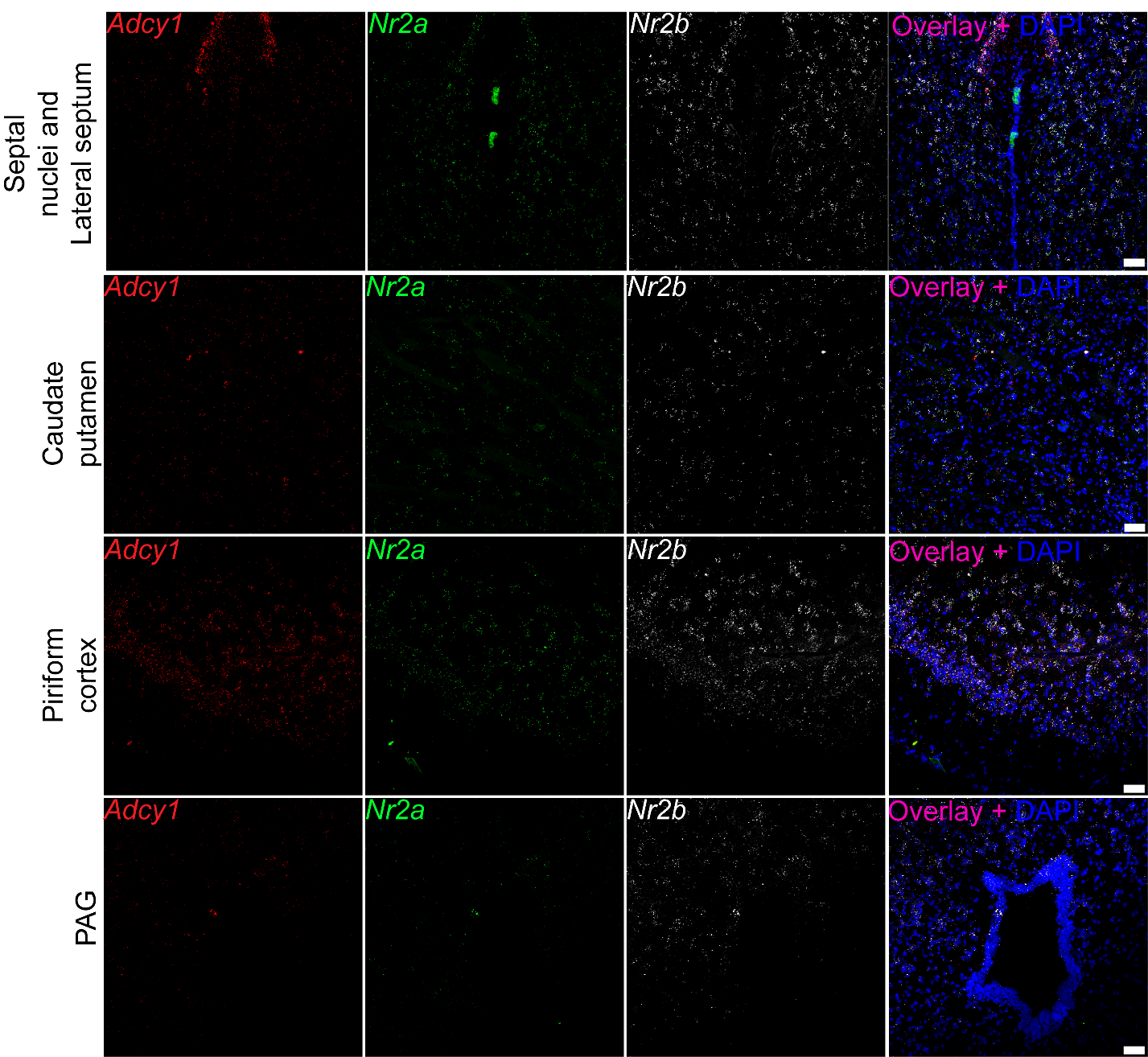

**Supplementary Figure 2. *Adcy1*, *Nr2a*, and *Nr2b* in the septal nuclei, lateral septum, caudate putamen, piriform cortex, periaqueductal gray (PAG), and spinal dorsal horn.** 20x; Scale bar = 50 µm.

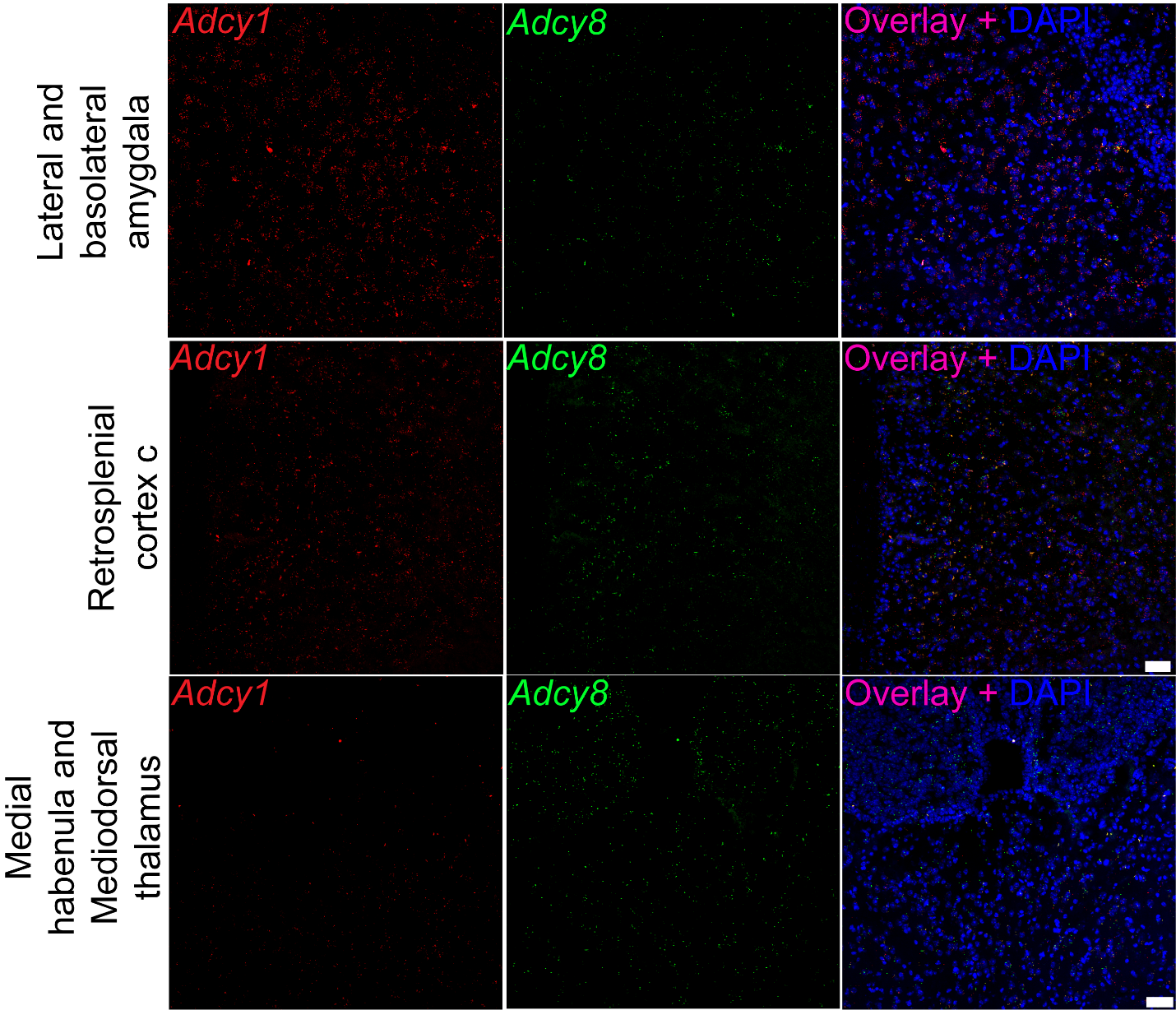

**Supplementary Figure 3. *Adcy1* and *Adcy8* in the lateral and basolateral amygdala, retrosplenial cortex c, medial habenula, mediodorsal thalamus.** 20x; Scale bar = 50 µm.

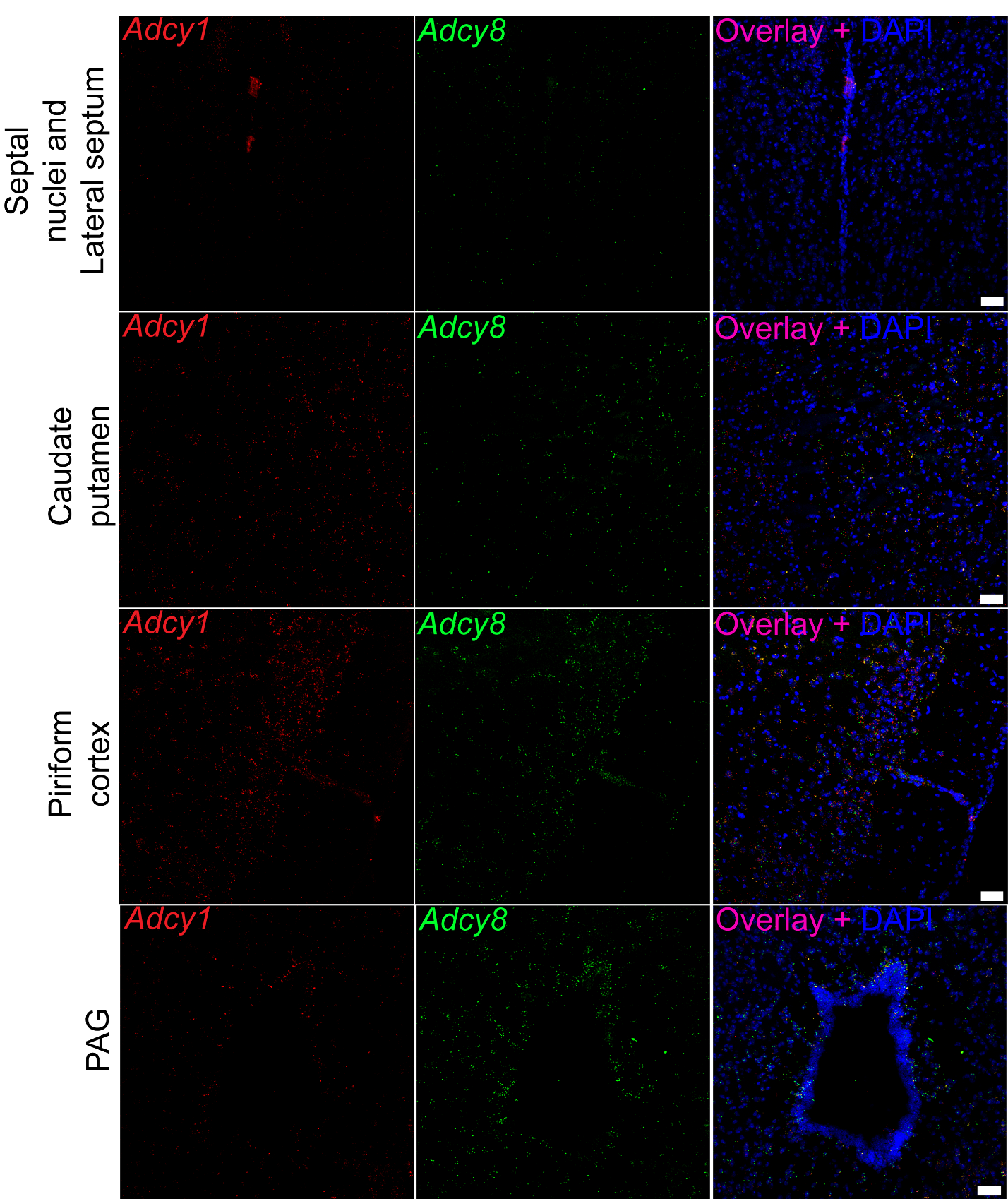

**Supplementary Figure 4. *Adcy1* and *Adcy8* in the septal nuclei, lateral septum, caudate putamen, piriform cortex, periaqueductal gray (PAG), and spinal dorsal horn.** 20x; Scale bar = 50 µm.

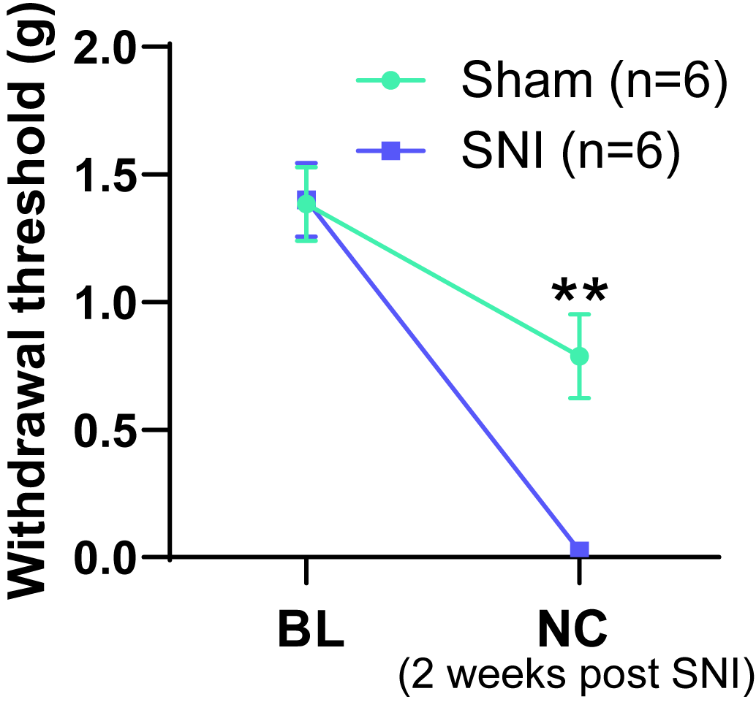

**Supplementary Figure 5. von Frey data for mice used in experiment 2 histology experiments.** Two-way ANOVA with Sidak multiple comparisons test **p<0.01
